# Fungal gasdermin-like proteins are controlled by proteolytic cleavage

**DOI:** 10.1101/2021.06.03.446900

**Authors:** Corinne Clavé, Witold Dyrka, Alexandra Granger-Farbos, Benoît Pinson, Sven J. Saupe, Asen Daskalov

**Author notes:** **Corresponding authors** Asen Daskalov, CNRS, UMR 5095, Institut de Biochimie et Génétique Cellulaires, University of Bordeaux, Bordeaux, France, Corinne Clavé, CNRS, UMR 5095, Institut de Biochimie et Génétique Cellulaires, University of Bordeaux, Bordeaux, France.

## Abstract

Gasdermins are a family of pore-forming proteins controlling an inflammatory cell death reaction in the mammalian immune system. The pore-forming ability of the gasdermin proteins is released by proteolytic cleavage with the removal of their inhibitory C-terminal domain. Recently, gasdermin-like proteins have been discovered in fungi and characterized as cell death-inducing toxins in the context of conspecific non-self discrimination (allorecognition). Although functional analogies have been established between mammalian and fungal gasdermins, the molecular pathways regulating gasdermin activity in fungi remain largely unknown. Here, we characterize a gasdermin-based cell death reaction, controlled by the *het-Q* allorecognition genes in the filamentous fungus *Podospora anserina*. We show that the cytotoxic activity of the HET-Q1 gasdermin is controlled by proteolysis. HET-Q1 loses a ∼5 kDa C-terminal fragment during the cell death reaction in presence of a subtilisin-like serine protease, termed HET-Q2. Mutational analyses and successful reconstitution of the cell death reaction in a heterologous host (*Saccharomyces cerevisiae*) suggest that HET-Q2 directly cleaves HET-Q1 to induce cell death. By analysing the genomic landscape of *het-Q1* homologs in fungi, we uncovered that the vast majority of the gasdermin genes are clustered with protease-encoding genes. These HET-Q2-like proteins carry either subtilisin-like or caspase-related proteases, which in some cases correspond to the N-terminal effector domain of NOD-like receptor proteins (NLRs). This study thus reveals the proteolytic regulation of gasdermins in fungi and establishes evolutionary parallels between fungal and mammalian gasdermin-dependent cell death pathways.

**Significance:** The recent discovery of gasdermin-like proteins in fungi have brought to light that this family of pore-forming proteins controls cell death in two of the major eukaryotic kingdoms, fungi and mammals. Yet, the regulation of cytotoxicity of the fungal gasdermins and their molecular pathways remain uncharacterized. Here, we describe the regulation through proteolytic cleavage of the fungal gasdermin HET-Q1 and uncover that majority of fungal gasdermins are genomically clustered with protease-encoding genes. Some of these genes encode proteins with caspase-related domains and/or are members of a family of immune receptors in mammals and plants. Overall, this work contributes towards our understanding of the evolution of gasdermin-dependent cell death, enlightening multiple evolutionary parallels between signaling pathways in mammals and fungi.

## Introduction

Spatiotemporally localized, regulated cell death (RCD) prevents cytoplasmic mixing and heterokaryon formation between conspecific individuals in fungi (1, 2). The cell death reaction limits the transmission of mycoviruses and other deleterious replicons, thus representing a defence reaction akin to an immune response (3, 4). The molecular characterization of cell death-pathways, regulating conspecific non-self discrimination (allorecognition) in fungi, has revealed evolutionary relations to cell death pathways operating in mammalian innate immunity, notably necroptosis and pyroptosis (5, 6). The extent of conservation of these pathways is gradually revealed as the characterization of fungal RCD pathways progresses.

Proteolytic cleavage is a general mechanism for regulation of programmed cell death (PCD) and RCD in metazoans and plants (7). In mammals, a family of cysteine-aspartic proteases (caspases) controls cellular suicide pathways like apoptosis and pyroptosis (8, 9). The latter represents a highly inflammatory, lytic cell death reaction, playing a central role in mammalian innate immunity (10, 11). Pyroptotic cell death is induced by pro-inflammatory caspases (caspase-1, -4, -5 and -11), which cleave a protein termed gasdermin D (GSDMD) (12, 13). The cleavage of GSDMD liberates the cytotoxic N-terminal domain of the protein (GSDMD-NT) from the inhibitory C-terminal domain (GSDMD-CT) and the processed GSDMD-NT disrupts the integrity of the plasma membrane by pore-formation (14–17). GSDMD belongs to a protein family comprising six members in humans (GSDMA, GSDMB, GSDMC, GSDME (also DFNA5) and PJVK (also DFNB59)) (18). Proteolytic cleavage has been shown to regulate the cytotoxic activity of other members of the gasdermin family, specifically of GSDME (19) and GSDMB (20). While the regulation of GSDME is caspase-3-dependent (19, 21), it has recently been reported that GSDMB is proteolytically activated by serine protease termed granzyme A (GZMA) (20).

Gasdermin-like proteins are abundant and widespread in fungi with some species encoding more than twenty gasdermin-homologs in their genomes (22). It has recently been established that a fungal gasdermin-like protein controls cell death in the context of allorecognition. The co-expression of antagonistic alleles of the *rcd-1* gene (regulator of cell death) triggers rapid, lytic cell death in asexual spores and hyphae of the model ascomycete *Neurospora crassa* (22). RCD-1 is a distant homolog of the cytotoxic pore-forming GSDMD-NT domain and, like gasdermin, induces cell death by targeting the plasma membrane, while forming oligomers (6). However, in spite of the uncovered functional similarities between the mammalian gasdermins and fungal RCD-gasdermin-like protein, in the latter case, cytotoxic activity is not dependent on proteolytic activation but results instead from the interaction of antagonistic RCD-variants.

Here, we characterized the fungal gasdermin *het-Q1* form *Podospora anserina* and report that its cytotoxic activity is regulated through proteolytic cleavage by a subtilisin-like serine protease named HET-Q2. *het-Q1* and *het-Q2* are idiomorphic genes (they represent alternate alleles of the same *het-Q* locus but are totally unrelated in sequence) and regulate cell death in the context of heterokaryon formation (somatic fusions between different strains). We demonstrated that a ∼5 kDa C-terminal fragment is cleaved from HET-Q1 in presence of HET-Q2. Expression of a truncated variant – HET-Q1(1-233) – was sufficient to reduce cell viability, while a point mutant – HET-Q1-F238A – preventing proteolytic cleavage of the fungal gasdermin, abolished its cytotoxicity. Catalytically inactive mutants of the HET-Q2 protease were equally unable to produce cell death when co-expressed with HET-Q1. The *het-Q1*/*het-Q2* RCD reaction can be reconstructed in the distant heterologous yeast host *Saccharomyces cerevisiae*, which strongly suggests that HET-Q2 removes directly the C-terminal inhibitory end of the gasdermin HET-Q1 to induce cell death. Importantly, *in silico* analyses of the genomic landscape of *het-Q1* homologs revealed that the fungal gasdermin genes are, in the vast majority of cases, the genomic neighbours of protease-encoding *het-Q2*-like genes. In addition, ∼20% of these protease-encoding neighbours carry a predicted caspase-like domain. Our work indicating that members of the gasdermin family in fungi are regulated through proteolytic cleavage reveals a remarkable evolutionary conservation of regulated cell death pathways in Eukaryotes.

## Results

### RCD during *het-Q*-allorecognition is controlled by idiomorphic genes

The *het-Q* locus is one of the last uncharacterized allorecognition loci in *P. anserina* (23). *P. anserina* strains either belong to the *het-Q1* or *het-Q2* genotype. Cellular fusions between strains from different *het-Q* genotypes are abortive and trigger RCD, which translates with the appearance of a ‘barrage’ (demarcation line) between such strains. The *het-Q* locus has been genetically located on chromosome 7 (24). To identify the precise genetic determinants controlling *het-Q* RCD, we analyzed the genetic distance between *het-Q* and the *PaATG1* locus (Pa_7_10890), situated on the same chromosome (25). We measured ∼2.5% of recombination between *het-Q* and *PaATG1*, which corresponds to ∼60 kbp in estimated physical distance between the two genes. We thus undertook a positional cloning approach with a DNA library of cosmids and plasmids from a *het-Q1* strain, covering a region of ∼100 kbp around *PaATG1* gene (*SI Appendix*, Fig. S1, Tables S1, S2, S3). Cosmid and plasmid clones were transformed into protoplasts of the incompatible *het-Q2* genetic background. Since co-expression of *het-Q1* and *het-Q2* is lethal, it is expected that clones bearing *het-Q1* show reduced transformation efficiency in this background (*SI Appendix*, Fig. S1; Table S1 and S2). Only constructs bearing the Pa_7_10775 gene showed reduced transformation efficiencies specifically in the *het-Q2* genetic background (*SI Appendix*, Fig. S1, Table S4). These experiments suggest that Pa_7_10775 is allelic to *het-Q1*. We replaced the Pa_7_10775 gene with a *nat1* cassette (nourseothricin resistance marker) producing a *ΔPa_7_10775* strain, which showed wild-type growth and fertility. The *ΔPa_7_10775* strain no longer produced a barrage reaction with a *het-Q2* strain, indicating that the RCD reaction is abolished (Fig. 1A). Ectopic re-introduction of Pa_7_10775 into the *ΔPa_7_10775* background restored the barrage reaction to *het-Q2* (Fig. 1A and 1B). These results indicate that Pa_7_10775 is *het-Q1*. The *ΔPa_7_10775* strain was thus designated *Δhet-Q*.

**Figure 1.**
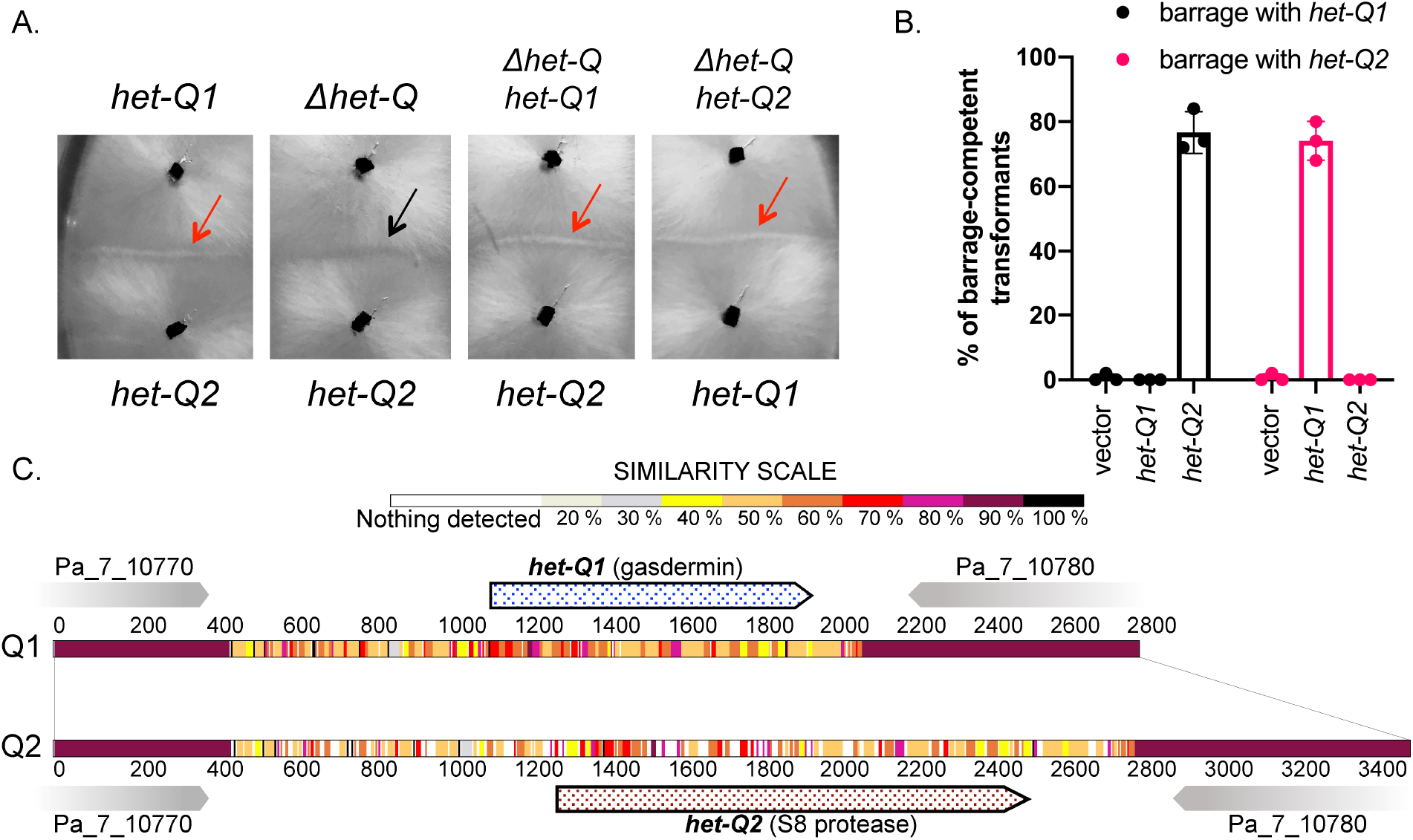
Gene idiomorphs *het-Q1* (gasdermin homolog) and *het-Q2* (S8 serine protease) define an RCD system in the context of heterokaryon incompatibility (allorecognition) in *Podospora anserina*. **A**. Cell death between *P. anserina* strains of antagonistic *het-Q1* and *het-Q2* genotypes manifest with the appearance of ‘barrage’ reaction between the fungal colonies (red arrows). Deletion of the *het-Q1* gene (Pa_7_10775) produces a *Δhet-Q* strain (we term it *Δhet-Q* as the *het-Q1* and *het-Q2* strains are isogenic with the exception of the *het-Q* locus) and abolishes the cell death reaction (barrage formation) with a *het-Q2* strain (black arrow). Reintroduction of *het-Q1* in a *Δhet-Q* strain restores the barrage reaction with a *het-Q2* tester strain, while introduction of *het-Q2* in the *Δhet-Q* background procures the ability to the transformants to produce a barrage reaction with a *het-Q1* tester strain. **B**. Quantification of *het-Q1*/*het-Q2* cell death, measuring the percentage of transformants producing barrage reaction with a *het-Q1* and a *het-Q2* tester strains. Experiments were performed in triplicate with 50 transformants tested per experiment. Empty vector has been used as a negative control. **C**. Graphic representation of the *het-Q* locus as found in strains from the *het-Q1* (Q1) and the *het-Q2* genotypes (Q2). Shown is an alignment of the *het-Q* locus from a *het-Q1* and *het-Q2* strain. Sequence similarity at nucleotide level is color-coded. Regions encoding the idiomorphic genes show low sequence similarity (in 40-60% range), while regions encoding the neighbouring genes (Pa_7_10770 and Pa_7_10780) show sequence similarity above 90% between the two genotypes.

Next, we amplified the same locus from a *het-Q2* strain using primers located in the ORFs flanking *het-Q1* (Pa_7_10775), Pa_7_10770 and Pa_7_10780 (*SI Appendix*, Fig. S1). The corresponding PCR fragment showed reduced transformation efficiency in the *het-Q1* background, while transformation efficiencies into the *het-Q2* or *Δhet-Q* backgrounds were at levels similar to that with the empty vector (*SI Appendix*, Fig. S1). *Δhet-Q* strains transformed with this PCR fragment acquired the *het-Q2* phenotype and triggered RCD with a *het-Q1* tester strain (Fig. 1). The *het-Q* locus situated between the Pa_7_10770 and Pa_7_10780 ORFs is totally dissimilar between the *het-Q1* and *het-Q2* strains and varies in length in the two backgrounds with 1.6 kbp and 2.2 kbp, respectively (Fig. 1C). The *het-Q1* and *het-Q2* genes are thus best described as idiomorphs rather than classical alleles (26).

### *het-Q1* encodes a gasdermin homolog and *het-Q2* a putative serine protease

The *het-Q1* gene encodes a 278 amino acid long protein, HET-Q1, which belongs to the same protein family as RCD-1 from *N. crassa* (6) and thus is a fungal gasdermin-like protein (*SI Appendix*, Fig. S2). We confirmed the homology between HET-Q1 and mammalian gasdermins, using the HHPred suite (27). The *in silico* analyses uncovered two regions of higher sequence conservation between HET-Q1, GSDMD-NT and GSDMA3-NT, detected around residues 148-155 and 210-217, as previously reported for RCD-1 protein (*SI Appendix*, Fig. S2).

The *het-Q2* ORF encodes a 366 amino acid long protein. HET-Q2 shows homology to subtilisin-like serine proteases of the S8 family (*SI Appendix*, Fig. S2). No signal peptide was detected in HET-Q2, which suggests that the predicted protease is not secreted. Based on sequence alignments and homology modeling, the amino acid residues D35, H105 and S266 of HET-Q2 were identified as constituting the catalytic triad of the putative protease. HET-Q2 lacks an N-terminal propeptide inhibitor domain (i.e. I9 domain), typically found in S8 subtilisin proteases (28).

### HET-Q2-dependent proteolytic cleavage of HET-Q1 triggers cell death

Considering the homology with mammalian gasdermins and the fact that these cell death-inducing proteins are activated by proteolytic cleavage, we reasoned that the HET-Q1/HET-Q2 cell death reaction might result from the proteolytic cleavage of the HET-Q1 gasdermin by the HET-Q2 serine protease. To test this hypothesis, we produced N-terminally V5-tagged HET-Q1 allelic variant (referred as V5-HET-Q1 or V5-Q1), which was functional and induced cell death at wild type levels (*SI Appendix*, Fig. S3). We then explored the proteolytic reaction by mixing crude cell extracts (cytoplasmic content) of strains expressing V5-tagged HET-Q1 with extracts from a *het-Q2* strain or from the *Δhet-Q* strain. Lysates mixtures (1:1 ratio) were incubated for up to 30 minutes (25°C). At the initial time point (at 0’ minutes) in both reaction mixtures (V5-Q1 + Q2 and V5-Q1 + ΔQ) the V5-tagged HET-Q1 protein was detected by Western blot at the expected molecular weight (∼35 kDa) (Fig. 2A). The molecular weight of HET-Q1 remained unchanged in the mixture of crude lysates from the *v5-het-Q1* and *Δhet-Q* strains (V5-Q1+ΔQ) (Fig. 2A). However, in presence of HET-Q2 (V5-Q1+Q2 mixture) we observed a progressive decrease in the amount of full-length HET-Q1 with a correlated appearance of a fragment of lower molecular weight (∼28-30 kDa), corresponding to the N-terminal part of the HET-Q1 protein (Fig. 2A). We concluded that in presence of the HET-Q2 serine protease, HET-Q1 is proteolytically processed and a ∼5 kDa C-terminal fragment (or domain) is cleaved from the protein.

**Figure 2.**
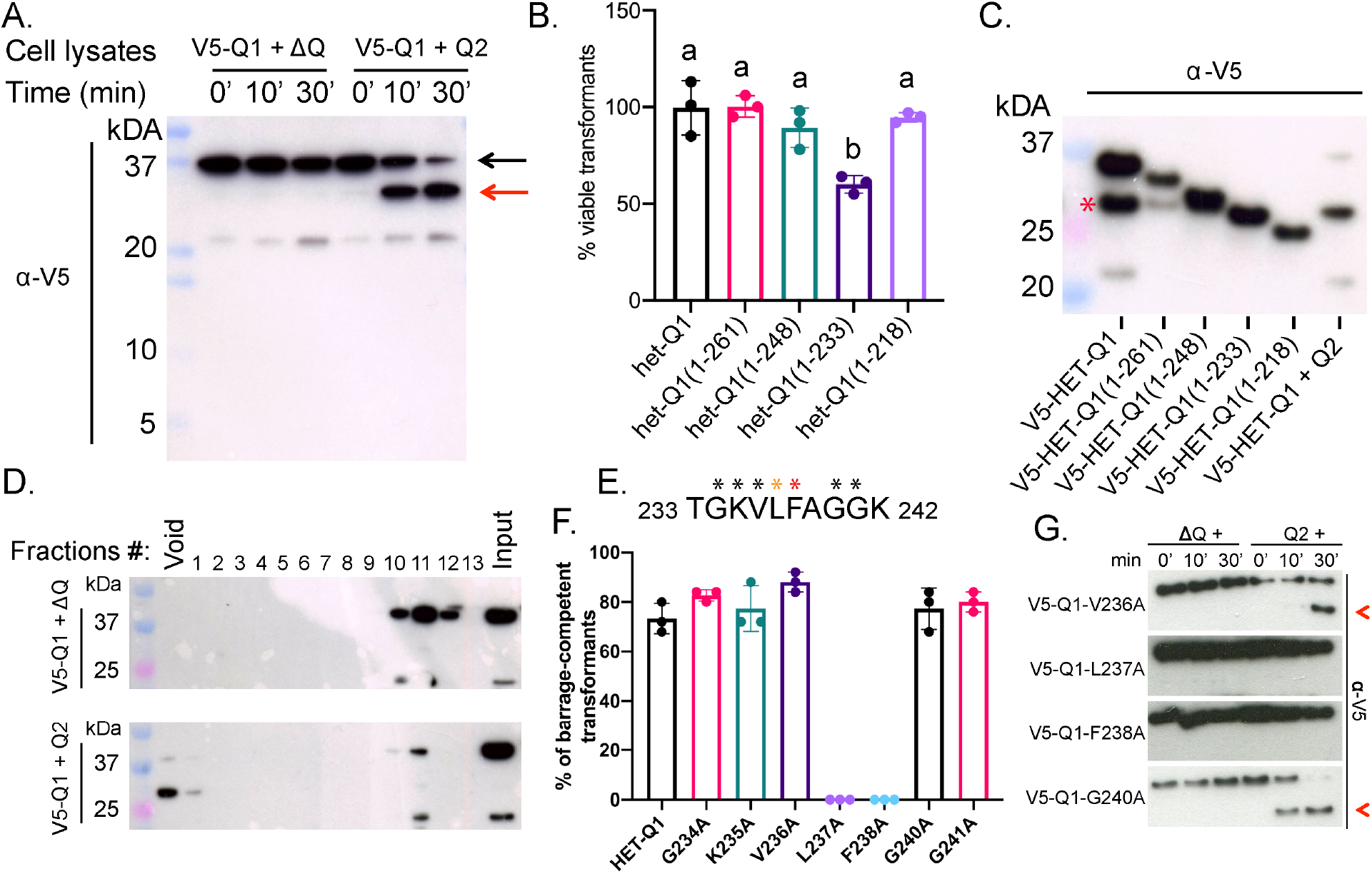
HET-Q2-dependent proteolytic cleavage of the gasdermin-like protein HET-Q1 induces cell death in *P. anserina*. **A**. Western blot showing accumulation of processed V5-HET-Q1 (V5-Q1) in a *het-Q2*-dependent manner. Crude cell lysates from V5-HET-Q1-expressing strain were mixed and incubated (for up to 30 min) with cell extracts from a *Δhet-Q* strain (V5-Q1 + ΔQ) or from a *het-Q2* strain (V5-Q1 + Q2). In the first reaction (V5-Q1 + ΔQ), V5-HET-Q1 remained unprocessed and appeared as single band at the expected size for the full-length protein (∼35-kDa) (black arrow). However, when mixed with extracts from a *het-Q2* strain (V5-Q1 + Q2), a smaller V5-HET-Q1 fragment (27-30-kDa) (red arrow) appears and accumulates in time, while the amount of full-length protein progressively decreases. **B**. Quantification of transformation viability of *Δhet-Q* transformants with truncated *het-Q1* alleles. Four truncations of *het-Q1* were introduced into *Δhet-Q* protoplasts and number of viable transformants counted. Three of the truncations (*het-Q1(1-261), het-Q1(1-248)* and *het-Q1(1-218)*) produced wild-type levels of viable transformants. Transformations with the fourth truncation – *het-Q1(1-233)* – resulted in ∼45% reduction of viable transformants. Experiments were performed in triplicate. P-value (a ≠ b) < 0.01, one-way ANOVA with Tukey’s multiple comparisons test. **C**. Western blot of full-length and truncated V5-HET-Q1 variants. Cytotoxic V5-HET-Q1(1-233) is of similar molecular size than processed V5-HET-Q1 in mixtures of cell extracts (V5-HET-Q1 + Q2). Protein bands corresponding to unspecific proteolysis due to high levels of protein expression are shown with red asterisk. **D**. Polyacrylamide gel electrophoresis (PAGE) and Western blot of FPLC (Fast protein liquid chromatography) fractions of crude cell extracts mixtures. Cell extracts containing V5-HET-Q1 were mixed with cell extracts from a *Δhet-Q* strain (V5-Q1 + ΔQ) (top panel) or from a *het-Q2* strain (V5-Q1 + Q2) (bottom panel). Processed V5-HET-Q1 in mixtures containing the HET-Q2 protease elutes in the void volume, containing protein aggregates of high molecular weight (above 1-MDa), while unprocessed V5-HET-Q1 elutes in fractions corresponding to proteins with lower molecular weight, close to that of monomeric V5-HET-Q1 (∼35-kDa). **E**. Protein sequence of HET-Q1(232-243). Amino acid residues substituted with alanine residues are marked with asterisks. Black asterisks indicate residues, which substitution did not impact the cell death reaction. Red asterisk indicates that the substitution abolishes the cell death reaction and yellow asterisk indicates that it impacts partially cell death. **F**. Quantification of barrage-inducing transformants expressing mutant V5-HET-Q1 variants. Experiments were performed in triplicate with 25 tested transformants per experiment. Mutant variants HET-Q1(L237A) and HET-Q1(F238A) are significantly affected in their cell death-inducing ability (P-value < 0.0001, one-way ANOVA with Tukey’s multiple comparisons test). **G**. Western blots of V5-HET-Q1 mutant variants in cell extract mixtures containing HET-Q2 protease (Q2 +) or without (ΔQ +). Mutations affecting the cell death reaction (V5-Q1-L237A and V5-Q1-F238A) prevent proteolytic cleavage of HET-Q1. Proteolytically processed mutants (V5-Q1-V236A and V5-Q1-G240A) are shown with red arrowheads.

To investigate whether the proteolytic cleavage activates the cytotoxicity of HET-Q1, we produced four truncated protein variants, removing short fragments in ∼15 amino acids increments from the C-terminal end of the protein. The truncations were introduced in the *Δhet-Q* strain and the number of viable transformants was quantified (Fig. 2B). Three of the tested truncations (HET-Q1(1-261), HET-Q1(1-248) and HET-Q1(1-218)) showed normal levels of transformation efficiency, while the fourth truncation – HET-Q1(1-233) – resulted in significant decrease in the number of viable transformants (Fig. 2B). Western blot analyses confirmed that the HET-Q1(1-233) truncation is of similar molecular weight to processed HET-Q1 (Fig. 2C). Noteworthy, among the truncation constructs, only strains expressing HET-Q1(1-261) produce barrage with a HET-Q2-expressing strain (*SI Appendix*, Fig. S3). These results suggest that some of the truncations impact the N-terminal cytotoxic domain of HET-Q1 and/or prevent the activation of the toxin by the protease. We concluded that the removal of the last 45 amino acids residues of HET-Q1 leads to an increased cytotoxicity of HET-Q1. In addition, we performed size-exclusion chromatography with cellular extracts. While unprocessed HET-Q1 elutes in low molecular weight fractions, we found that in mixtures containing HET-Q1 and HET-Q2 (Q1+Q2) proteolytically processed HET-Q1 elutes in the void fraction. This result suggests that the proteolytic cleavage induces a transition from monomeric towards multimeric state of the fungal toxin, similarly to members of the gasdermin family in mammals (Fig. 2D) (18).

Next, we undertook to identify amino acid residues important for the proteolytic cleavage of HET-Q1 and replaced individual residues with alanines in the region where the cleavage occurs. We generated a set of seven HET-Q1 mutants, encompassing the region from residues G234 to G241 of HET-Q1 (Fig. 2E) and tested the ability of the mutants to induce cell death with a strain expressing HET-Q2 (Fig. 2F). The majority of HET-Q1 mutants were not affected in their ability to induce cell death. Only strains expressing HET-Q1-L237A and HET-Q1-F238A were unable to produce barrage with the *het-Q2* strain (Fig. 2F). However, after prolonged contact between transformants expressing HET-Q1-L237A and the *het-Q2* strain, we observed the formation of attenuated barrages, suggesting that the mutation affects the cell death reaction only partially. We then probed whether the proteolytic cleavage of the HET-Q1-L237A and HET-Q1-F238A mutants occurs in presence of HET-Q2 (Fig. 2G). Crude cellular extracts from transformants expressing HET-Q1 mutant variants were mixed in a 1:1 ratio with cellular extracts from a *Δhet-Q* strain or *het-Q2* strain and incubated for up to 30 minutes. Mutations affecting the cell death reaction (L237A and F238A) were found to equally affect the proteolytic cleavage of HET-Q1. The two mutant variants (HET-Q1-L237A and HET-Q1-F238A) remained unprocessed even after 30 minutes in presence of HET-Q2 (Fig. 2G). Mutant HET-Q1 variants that did not affect the cell death reaction (V236A and G240A) did not abolish the proteolytic cleavage of HET-Q1 and were processed after 30 minutes in presence of HET-Q2 (Fig. 2G). We concluded from these experiments that the amino acid residues L237 and F238 are important for the proteolytic cleavage of HET-Q1 and likely integral to the cleavage site. The results further confirm that proteolysis of HET-Q1 is necessary for the induction of cytotoxicity.

### Proteolytic activity of the subtilisin-like protease HET-Q2 is essential for gasdermin cytotoxicity

We reasoned that HET-Q2 might be constitutively active (or alternatively be specifically activated by the interaction with HET-Q1) and that its proteolytic activity controls the cell death reaction, likely by direct cleavage of HET-Q1. We therefore generated three *het-Q2* mutants, replacing each of the residues forming the catalytic triad of HET-Q2 (D35, H105 and S266) with an alanine amino acid residue. The mutant alleles (*het-Q2-D35A, het-Q2-H105A* and *het-Q2-S226A*) were introduced into *Δhet-Q* background and transformants were tested for barrage formation with the *het-Q1* tester strain. While ∼76% of transformants with wild-type *het-Q2* produced a barrage reaction with the *het-Q1* strain, transformants with mutant *het-Q2* alleles showed reduced or abolished cell death-inducing activity (Fig. 3A). Strains transformed with two of the mutants – *het-Q2-H105A* and *het-Q2-S226A –* were unable to trigger cell death with a *het-Q1* strain, while only ∼37% of transformants with the *het-Q2-D35A* allele produced a barrage reaction (Fig. 3A). The decrease of activity shown by *het-Q2-D35A* was statistically significant and consistent with previously reported residual protease activity for an equivalent mutation in a human subtilisin (29). These results demonstrate that the proteolytic activity of HET-Q2 is necessary for the control of the HET-Q1-dependent cell death reaction.

**Figure 3.**
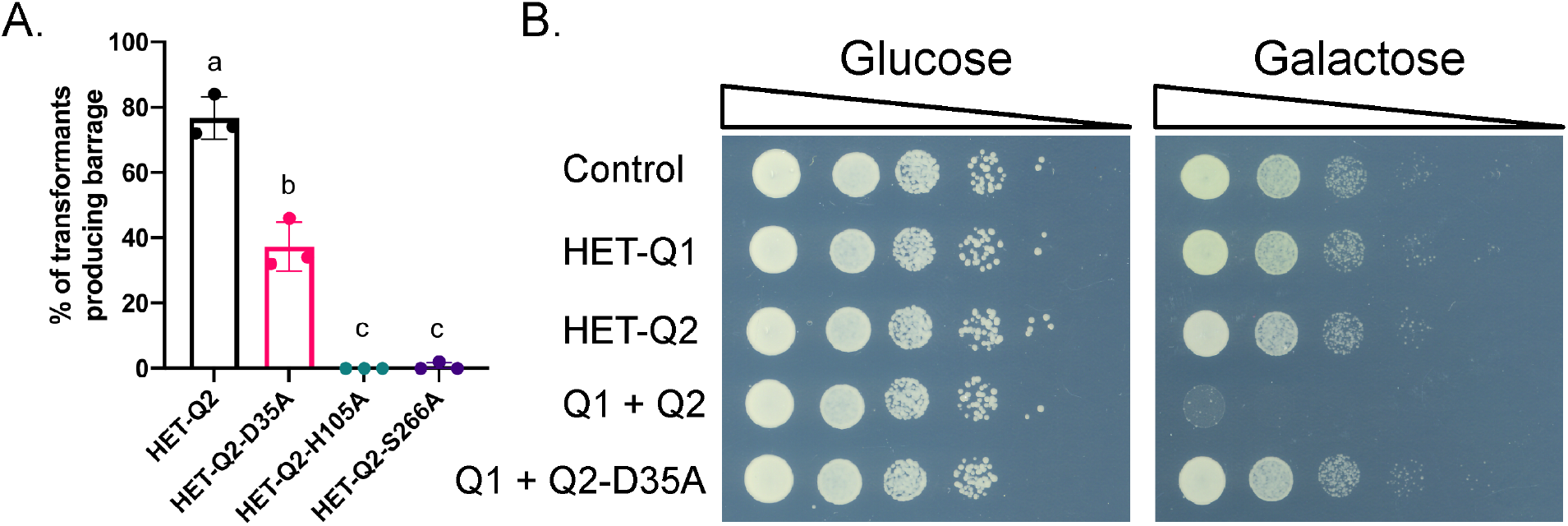
Proteolytic activity of the subtilisin-like protease HET-Q2 is essential for gasdermin cytotoxicity. **A**. Quantification of cell death (barrage) induction by transformants expressing HET-Q2 and mutant HET-Q2 variants. Alanine substitutions of residues forming the predicted catalytic triad of HET-Q2 (D35A, H105A and S266A) affect the cell death reaction. Experiments were performed in triplicate with 50 transformants per experiment. P-value (a ≠ b ≠ c) < 0.0001, one-way ANOVA with Tukey’s multiple comparisons test. **B**. *Saccharomyces cerevisiae* spot test essay on repressed (glucose) and inducible (galactose) growth media. Co-expression of HET-Q1 with HET-Q2 (Q1 + Q2) leads to growth arrest of *S. cerevisiae*, while individual expression of both proteins or co-expression of the gasdermin with a catalytically affected protease variant (HET-Q2-D35A) did not affect yeast growth.

Next, we decided to use *Saccharomyces cerevisiae* as a heterologous system, in which to reconstitute the HET-Q1/HET-Q2 cell death reaction. We reasoned that cell death signaling pathways between the yeast and *Podospora* are unlikely to be conserved, as the two fungal species are phylogenetically very distant and differ in cellular organization (unicellular versus multicellular). None of the known components of *P. anserina* allorecognition pathways has homologs in yeast. We co-expressed HET-Q1 and HET-Q2 in yeast. The *het-Q1* gene was under the control of a constitutive promoter, while *het-Q2* was under a galactose-inducible *gal1* promoter. Experiments were performed at 30°C, the optimal growth temperatures for *Saccharomyces*. We found that co-expression of HET-Q1 and HET-Q2 strongly affected the growth of *S. cerevisiae* (Fig. 3B). Expression of HET-Q1 alone or HET-Q2 alone did not alter the growth of *S. cerevisiae* nor did the HET-Q2-D35A mutant when co-expressed with HET-Q1 (Fig. 3). Taken together, these results suggest that *het-Q1* and *het-Q2* genes constitute an autonomous cell death induction system and further support the direct cleavage of the HET-Q1 gasdermin by the HET-Q2 serine protease.

### Genomic clustering of *het-Q1* homologs with protease-encoding *het-Q2-like* genes in fungi

Gasdermin-encoding *het-Q1* homologs are ubiquitous in Ascomycota and have been identified in more than 180 fungal species (6, 22). The uncovered functional relation between HET-Q1 and the HET-Q2 protease, as well as the previously reported genomic clustering of regulation and cell death execution modules in fungal necroptosis-like pathways (5, 30), prompted us to analyze the local genomic landscape of *het-Q1* homologs in fungi. We retrieved 1884 *het-Q1* homologs situated in the genomes of 401 fungal species using *P. anserina het-Q1* as a query sequence. We found that 80% of the *het-Q1* genes had at least one gene encoding for a protein with a protease domain as close neighbor (within 10-kbp) (Fig. 4A and 4B, *SI Appendix*, Table S5). Such genes were frequently adjacent to the gasdermin-encoding *het-Q1* homologs – forming two-gene clusters – and were termed *het-Q2*-like (Fig. 4A). Approximately 63% (1187) of the analyzed 1884 fungal gasdermins were clustered with *het-Q2*-like genes encoding proteins with a putative S8 serine protease domain, as found in HET-Q2 (Fig. 4B). Remarkably, another 19% (353) of the *het-Q1* homologs were in the vicinity of a *het-Q2*-like gene encoding for a protein with a predicted protease domain of the CHAT (Caspase HetF Associated with TPRs) clade, which belongs to the same protease family as the mammalian caspases (31). We also found that 20% (373) of the *het-Q1* homologs were not situated near a *het-Q2*-like gene, yet the vast majority of *het-Q1* genes in this category (76%) were from fungal genomes containing at least one *het-Q1*/*het-Q2*-like two-gene cluster (*SI Appendix*, Table S5). Overall, we identified 1617 *het-Q1*/*het-Q2*-like clusters (1417*het-Q1/het-Q2-*like gene pairs and 88 *het-Q1*/two *het-Q2-*like three-gene clusters) from 366 fungal species with a dozen of species carrying at least ten such clusters in their genomes (*SI Appendix*, Table S5). Noteworthy, we also found two distant *het-Q1* homologs (Pa_6_6615 and Pa_5_3510) in the genome of *P. anserina*, clustered with *het-Q2*-like genes, one being of the S8 type, the other of the CHAT-type (*SI Appendix*, Fig. S4, Table S5).

**Figure 4.**
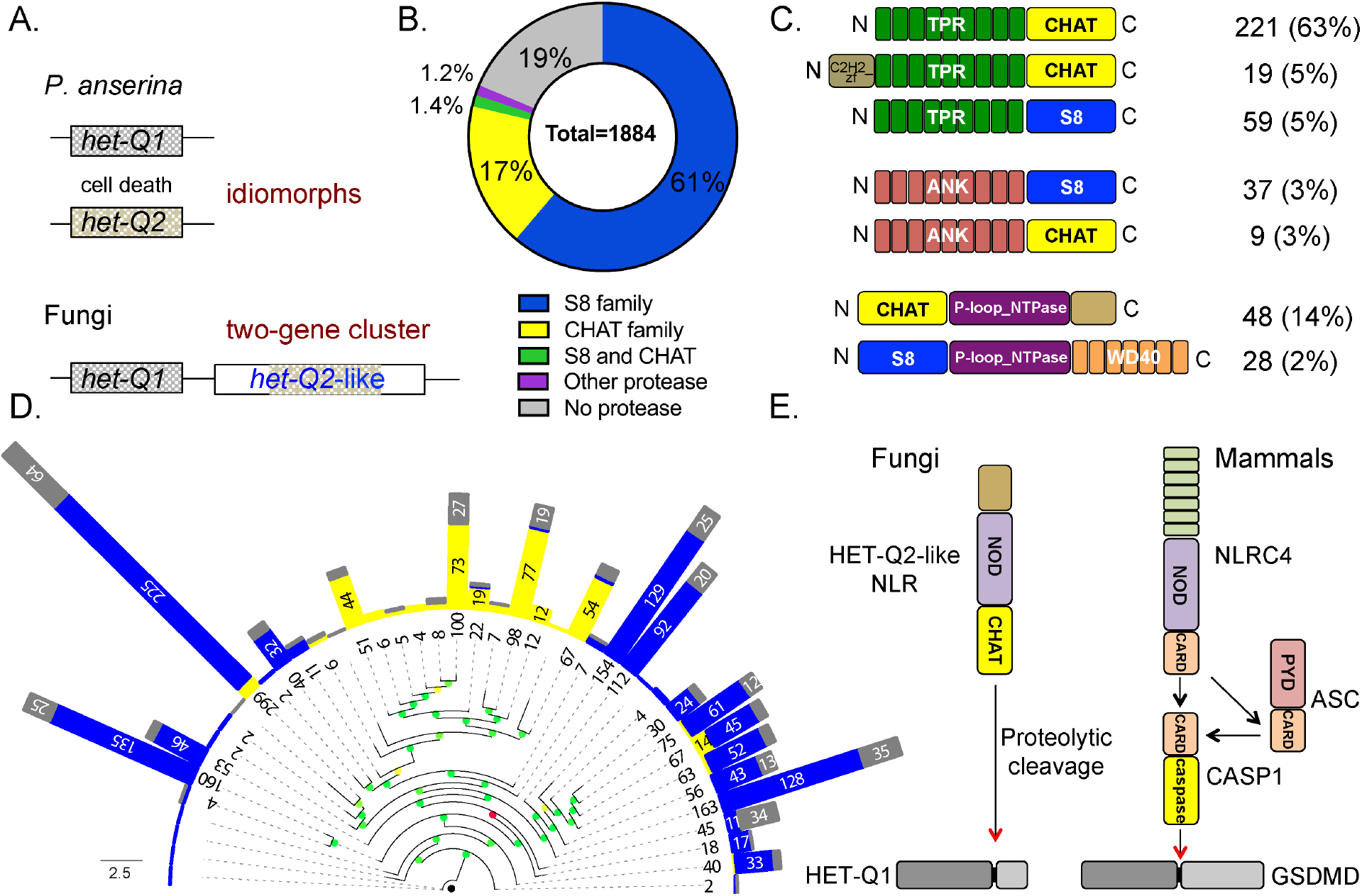
Gasdermin *het-Q1* homologs are genomically clustered with protease-encoding *het-Q2*-like genes in filamentous fungi. **A**. Cartoon representation of *het-Q1*/*het-Q2* gene idiomorphs, defining the allorecognition cell death reaction in *P. anserina* and of *het-Q1*/*het-Q2*-like gene clusters, distributed broadly in Fungi. **B**. Percentage of gasdermin homologs situated in the vicinity (10 kbp upstream and downstream) of protease-encoding *het-Q2-*like genes. **C**. Cartoons representing annotated HET-Q2-like protein architectures. Given are the numbers of proteins in each category and the percentage that each category represents from the total number of CHAT- or S8-containing HET-Q2-like proteins. **D**. Maximum-likelihood phylogenetic tree of *het-Q1* genes. Numbers of *het-Q1* homologs in collapsed branches are shown for each branch. Branch support is color-coded: red – 0.00, yellow – 0.50, grass-like green – 0.75, green – 1. Coloured bars at tip of branches show the type of protease domain found on the clustered HET-Q2-like proteins (blue – S8 family, yellow – caspase-like CHAT family) or the number of het-Q1 homologs not clustered with a *het-Q2*-like gene (grey). **E**. Comparative models of cell death pathways in Fungi and Mammals. Shown is an example of a CHAT-domain containing fungal NLR-like protein – a HET-Q2-like protein – hypothesized to activate through proteolytic cleavage a downstream HET-Q1 protein. Such HET-Q2-like/HET-Q1 pathways appear analogous to pyroptotic-inducing cell death pathways in mammals. Given example shows the downstream activation of GSDMD (gasdermin D) by NLRC4 (NLR family CARD-containing protein), which can directly activate caspase-1 (CASP1) or indirectly via the ASC protein (Apoptosis-associated speck-like protein containing a CARD) (40). NOD – Nucleotide-binding and oligomerization domain; CARD – Caspase-activation and recruitment domain; PYD – Pyrin domain.

The fungal gasdermins show a considerable phylogenetic diversity and formed more than 20 well-supported clades (support values above 0.9 for majority of non-singleton branches) comprising between ten and 300 *het-Q1* members (Fig. 4D). Although exception occurred, each clade or super-clade was predominantly associated with either the S8 and CHAT-protease type (Fig. 4D). This observation underscores further the functional link between the clustered genes and suggests that gasdermin and protease encoding-genes spend considerable evolutionary time in association while occasional re-association with a distinct protease-type can occur.

To gain insight into the functional role of these cell death pathways, we analyzed the domain architectures of the HET-Q2-like proteins (Fig. 4C, *SI Appendix*, Table S6). In the set of 1601 HET-Q2-like protein sequences, 1435 (90%) carry N-(1374) or C-termini (125) longer than 100 amino acids, in addition to the S8/CHAT protease. Using Pfam profiles, we annotated 493 of the 1499 long N- and C-termini (33%). The vast majority of the annotated sequences (above 95%) belonged to 16 Pfam clans and resulted in 18 potential protein architectures of HET-Q2-like proteins (Fig. 4C, *SI Appendix*, Table S6). Several of the most abundant HET-Q2-like protein architectures presented annotated domains consisting of tandem-repeat motifs (TPR, ANK or WD40 repeats), which are known to mediate protein-protein interactions and/or ligand binding functions (32) (Fig. 4C). Furthermore, we found that some of the *het-Q2*-like genes encode for NOD-like receptors (NLRs), exhibiting a typical tripartite domain organization with a central P-loop containing NOD (Nucleotide-binding and oligomerization) domain flanked by the protease domain and tandem-repeat motifs (specifically S8 / NACHT / WD40) (Fig. 4C). Overall, our analyses indicate that the majority of fungal gasdermins are clustered with genes encoding for putative CHAT or S8 protease domain-containing, multi-domain proteins, some of which belong to a class of cell death-controlling innate immunity receptors.

Based on these findings, we postulate that fungal gasdermins are in general regulated by proteolytic cleavage by HET-Q2-like functional partners (encoded by a closely linked gene) and are an essential part of broader, diverse and widespread signaling pathways in fungi (Fig. 4E). The genomic clustering of the gasdermin genes and the gene encoding their putative cognate protease is highly reminiscent of the clustering of HeLo-domain cell death execution proteins with their cognate regulatory amyloid signaling NLR (30, 33). In this context it is noteworthy, that the number of gasdermin homologs and Helo-domain pore-forming toxin homologs in fungal genomes are correlated (Pearson’s r=0.65, p-value < 1e-55, Spearman’s ρ=0.49, p-value < 1e-27) (*SI Appendix*, Table S7), suggesting that the life-style of certain species calls for a high number of parallel RCD pathways.

## Discussion

In this study, by molecularly characterizing a fungal allorecognition system from *Podospora anserina*, we show that fungal gasdermin-like proteins are controlled by proteolytic cleavage, similar to their mammalian counterparts (13, 14). We uncovered that the fungal gasdermin HET-Q1 is proteolytically processed in the presence of the serine protease HET-Q2, during the allorecognition process in *P. anserina*. The proteolytic cleavage removes a ∼5 kDa C-terminal fragment, which unleashes the cytotoxic activity of HET-Q1. We reconstituted the cell death reaction by co-expressing HET-Q1 and HET-Q2 in the yeast *Saccharomyces cerevisiae*. The latter finding strongly suggests that the fungal gasdermin is directly processed by the serine protease and establishes HET-Q1/HET-Q2 protein pair as an autonomous RCD system. In this context, we uncovered by genome mining that ∼80% of the gasdermin-encoding genes in fungi are clustered with *het-Q2*-like genes, encoding proteins with a putative protease domain, which belongs in the majority of cases to the subtilisin-like serine proteases (S8 family). Our analyses also identified that a subset of the HET-Q2-like proteins carry a caspase-related CHAT domain, while others exhibit NLR-like protein architectures. These findings prompted us to propose that similarly to HET-Q1, proteolytic cleavage is the general mode of activation for the fungal gasdermins. Furthermore, we propose that for the vast majority of HET-Q1 homologs the proteolysis occurs via an activated HET-Q2-like protein, encoded by the gene adjacent to the gasdermin-encoding gene. Genomic clustering of genes encoding pore-forming toxins and the receptors that activate them has been previously reported in fungi in the case of the HeLo-domain pore forming toxins and their cognate NLR receptors (30, 33).

### Gasdermin-regulated cell death in the context of comparative immunology

This study extends the evolutionary parallels between fungal and mammalian cell death pathways beyond the previously uncovered conservation of cell death execution modules (gasdermin pore-forming domain). Our findings establish a broader evolutionary framework for gasdermin-regulated cell death by unveiling a common mode of activation (proteolysis) for the gasdermin family in Eukaryotes, performed by similar type of proteases, which can be associated – at least in some cases – to homologous upstream receptors (NLR proteins) (Fig. 4E).

During pyroptosis the mammalian gasdermins are processed either by caspases (11–13, 34) or in the case of GSDMB by a serine protease (GZMA) (20). Serine proteases play a key role in human immunity, where they can control inflammatory and non-inflammatory cell death pathways (35). Furthermore, subtilisin-like serine proteases termed *phytaspases* (plant aspartate-specific protease) have been shown to control cell death in plants (36, 37). Phytaspases exhibit caspase-like specificity and can induce immune-related cell death in response to viral infections (38). Our findings now suggest that both types of proteases (subtilisin-like and caspase-like) regulate gasdermin-dependent cell death in fungi, likely in the context of organismal defense. While metacaspases (39), distantly related to caspases, have been previously shown to control fungal cell death in various physiological and stress conditions (1), our work proposes an unprecedented role for caspase-like domains in fungal non-self recognition processes.

Beside the significance for the evolution of proteolytic control of RCD, our work uncovers that, as described in mammals, some gasdermin proteins are downstream of NOD-like receptors or NLR-like proteins. Nevertheless, it appears that in fungi the signaling pathways leading to activation of gasdermin cytotoxicity are less complex with the NLR protein carrying directly the protease domain, while in mammals the NLRs use adapter domains (i.e. CARD – Caspase activation and recruitment domain) to activate downstream caspases directly (in some cases) or though adaptor proteins like ASC (Apoptosis-associated speck-like protein containing a CARD) (Fig. 4E) (40).

### Allorecognition systems in fungi: branching out of broader and unexplored cell death pathways

Our findings suggest that the *het-Q1*/*het-Q2* allorecognition system found in *P. anserina* has emerged from an ancestral *het-Q1*/*het-Q2-*like two-gene cluster. In this model, genomic rearrangements have resulted in the *het-Q2* gene, which encodes only the protease domain of a HET-Q2-like protein and hence results in a constitutively active protease, triggering cell death when co-expressed with HET-Q1. In this hypothesis, the *het-Q1/het-Q2* system would represent an evolutionary exaptation by which a preexisting immune pathway has been refurbished into an allorecognition system (41). A similar evolutionary scenario has been proposed for a two gene cluster involving a NLR and a different pore-forming toxin (HET-S) in *Podospora*, where a point mutation and a transposon insertion have generated an incompatibility system (*het-s*/*het-S* allelic system) (42). In addition, several other NLR-based allorecognition systems are considered to originate from broader non-self recognition-dedicated molecular pathways (43). The exaptation model could equally be proposed for the *rcd-1-1*/*rcd-1-2* allorecognition system in *N. crassa*, where genomic rearrangements have been associated with the *rcd-1* locus (22). Our study supports thus further the hypothesis that allorecognition genes originate from pre-existing broader signaling pathways, likely involved in xenorecognition and mediation of interspecific biotic interactions (44).

### Conclusion

The present results establish that in filamentous fungi, in addition to necroptotic-like pathways regulated by amyloid signaling (5), gasdermin-dependent pathways regulated by proteolytic cleavage abound. Mammals and fungi thus share core components and molecular mechanisms of RCD pathways. We hereby critically expand the evolutionary parallels between RCD pathways in two of the major clades of Opisthokonta (fungi and animals). While the extent of the molecular and functional similarities (and differences) remain to be fully explored, our study unveils a remarkable evolutionary conservation in cell death signaling and execution in the context of eukaryotic innate immunity and as such represents a further step towards the molecular definition of the fungal immune system.

## Methods and materials

### Strains and plasmids

*Podospora anserina* strains used in this study were wild type *het-Q1* and *het-Q2* and the *Δhet-Q1* (*Δhet-Q*) strain. All strains were cultivated at 26°C using standard procedures. To produce the *Δhet-Q* strain, we replaced the *het-Q1* ORF (Pa_7_10775) with the *nat1* gene, encoding a nourseothricin acetyl transferase, procuring resistance to the nourseothricin antibiotic. The *het-Q1* deletion cassette consists of the *nat1* gene flanked by the upstream and downstream regions of the *het-Q1* gene. To produce the deletion cassette, we amplified 1055 bp upstream of *het-Q1* ORF using primers 5’ AATATTGGAGAAGCAAAATGAGGC 3’ and 5’ TTTAGGTTCTTCTGAAAAGAGATAC 3’ and cloned the PCR fragment upstream of the *nat1* gene. Downstream of the *nat1* gene, we cloned a PCR fragment corresponding to 894 bp downstream of Pa_7_10775, starting from the stop codon of the gene. The PCR fragment was amplified with the primers 5’ GAGGAAGGTTATAATCGATGTGTG 3’ and 5’ GCAGTGGCGTTTGTCTTCGC 3’. The *het-Q1* deletion cassette was constructed into a pBlueScript II-derived vector (termed ‘p1’). To delete the *het-Q1* gene, we used 5 µg of 2.76 kbp *NcoI*-linearized 10775::nat1 (deletion cassette) DNA fragment, which we introduced into a *het-Q1 ΔPaKu70* strain. Nourseothricin-resistant transformants were selected, PCR screened and backcrossed with the wild type reference strain to obtain *Δhet-Q* homokaryons. The barrage phenotype segregated with the nourseothricin-resistance phenotype in the progeny with resistant transformants unable to produce a barrage reaction with a *het-Q2* strain.

The *het-Q1* gene was amplified with primers 5’ GTGTTTCGCTTGTTCAATCCG 3’ and 5’ TTTTCACAATGGAAATCGGGATG 3’ and the 1.85 kbp PCR fragment was cloned with *EcoRV* into a pBlueScript-derived vector carrying the hygromycin resistance *hph* gene. Similarly, we amplified the *het-Q2* gene using primers 5’ ACGATCAAACAACCCAGTTCGG 3’ and 5’ GAACCCCGATTTCAAGTATATGC 3’ and the PCR fragment (2.55 kbp) was cloned with the *EcoRV* restriction enzyme into the SKhph vector. The final constructs were sequenced and the vectors (SKhph, SKhph-Q1, SKhph-Q2) were used for viability reduction essays.

For heterologous expression of *het-Q1* and *het-Q2* in the yeast *Saccharomyces cerevisiae*, the CDS (coding sequences) of the two genes were PCR amplified and cloned in the pRS413 (45) and pGal vectors, respectively. The *het-Q1* sequence was amplified with primers 5’ TTGGATCCATGCCCACCAAAACCTCCCAAC 3’ and 5’ GGTCGACTCACGCCTTGGCAGCAACATC 3’, carrying BamHI and SalI restriction sites. The 853 bp PCR fragment was cloned in the pRS413 vector digested with *BamHI* and *XhoI*, to obtain the pRS413-het-Q1 plasmid. The *het-Q1* sequence was under the control of the strong constitutive promoter of the *gpd* (glyceraldehyde-3-phosphate dehydrogenase) gene. The *het-Q2* coding sequence was cloned under the control of a galactose-inducible promoter (*gal1*), using ‘gap repair’ methods (46), to produce the pGal-het-Q2 plasmid.

Site-directed mutagenesis was performed with QuikChange II kit (Agilent), using manufacturer recommended procedures.

### Mapping and positional cloning of *het-Q1* and *het-Q2*

To identify the *het-Q* locus and clone the *het-Q1* and *het-Q2* genes, we first undertook a classical genetics mapping to situate more precisely the locus on chromosome 7. For this we crossed a *P. anserina het-Q2* strain with a *het-Q1 ΔPaATG1* strain (*PaATG1::hph)*, where the *PaATG1* is replaced with hygromycin resistance cassette. The *PaATG1* gene deletion has been previously described and characterized (25). We isolated 112 uninucleated spores, from which 78 spores germinated. Hygromycin-resistance screen showed that 35 (45%) spores are resistant to the antibiotic, while 43 (55%) are sensitive. Barrage tests (with *het-Q1* and *het-Q2* tester strains) further showed that only 2 recombinant spores (*het-Q2 ΔPaATG1)* have been obtained from the 78 germinated spores. Based on the 2.5% of recombination between the *PaATG1* and the *het-Q1* loci, and knowing that 1 cM equals ∼25 kbp in *P. anserina*, we calculated that the physical distance between the two loci is ∼65 kbp.

To pinpoint the *het-Q* locus, we undertook a ‘chromosomal walking’ approach (47) with cosmids and plasmids carrying DNA from the *het-Q1* strain, previously used to sequence *P. anserina* reference genome (S genotype) (48). We used four cosmids (GA0AAD19ZF04 (CQ2), GA0AAD26ZD03 (CQ3), GA0AAD29ZA05 (CQ4) and GA0AAD50ZA08 (CQ5)) and five plasmids (GA0AB158BG07 (Q1), GA0AB263AH07 (Q2), GA0AB26BB11 (Q3), GA0AB218BA11 (Q4) and GA0AB91DC06 (Q5)) encompassing a 200-kbp region around the *PaATG1* (Pa_7_10890) locus. Cosmids were carrying the *hph* gene (hygromycin resistance) and were transformed in a *het-Q2* strain and viable transformants quantified. Transformations were performed with 10 µg of cosmid DNA. Plasmids used for the identification of the *het-Q* locus were used in co-transformation with the *hph*-bearing vector pCSN43 (GenBank: LT726869.1) (3:1 ratio for 12 µg total DNA). These viability reduction assays identified the plasmid GA0AAD29ZA05 (CQ4), carrying an 11-kbp insert and containing 5 genes, as the region, in which *het-Q1* is situated. Five variable in size deletions were produced and after viability reduction tests, it was determined that vectors carrying DNA fragments containing the Pa_7_10775 gene reduced the number of hygromycin-resistant transformants.

### Viability reduction and vegetative incompatibility essays

Vegetative incompatibility essays (barrage tests) were performed on standard corn meal agar DO medium. Viability essays were performed by transformation of protoplasts (or spheroplasts) of *P. anserina* strains (49)(of the *het-Q1, het-Q2* or *Δhet-Q* genotypes) with the SKhph, SKhph-Q1 and SKhph-Q2 vectors carrying the *hph* gene, which procures hygromycin resistance. Transformations were carried with 7.5 or 10µg of vector DNA. Transformants were counted after 6 days of incubation at 26°C.

### *Saccharomyces cerevisiae* spot test essay

The pGal-het-Q2, pRS413-het-Q1 and the two vectors (pGal and pRS413) expressing free GFP (Green Fluorescent Protein) were transformed or co-transformed in a *S. cerevisiae* (mat-a, his3Δ1, leu2Δ0, ura3Δ0, trp1::LEU2) using the One-step transformation protocol (50). Tryptophan and histidine prototrophic transformants were selected and grown in liquid media (10% glucose). Culture suspensions (1.5.10^7^ cellules/ml) were serially diluted (by a factor of 10) and 5 µl of each suspension were spotted on synthetic media with galactose (2%) or glucose (2%) as carbon source. Transformants were incubated at 30°C from 3 to 5 days.

### Crude cell extracts and *in vitro* proteolytic cleavage of HET-Q1

Protoplasts of different *P. anserina* strains were prepared using standard methods (51) at a final concentration of 10^6^ protoplasts/µl. Cell extracts were prepared from 200 µl of protoplasts, which were suspended in 10 mM phosphate buffer (Na_2_HPO_4_, NaH_2_PO_4_) pH 7.4 after centrifugation at 4°C, 5000 rpm for 10 min. Protoplast suspensions were sonicated (at amplitude four with a Soniprep 150) twice for 10s with 30s on ice between sonication cycles. Sonicated samples were centrifuged at 4°C, 10 000 rpm for 10 min and 160 µl of supernatant were transferred to a new Eppendorf tube and kept at 4°C.

Experiments were carried by mixing 50 µl of each crude extract and incubating the reaction mixtures at 26°C up to an hour. To stop the reaction at a specific time-point, 15 µl of the crude cell extract mixtures were added to 15 µl of 3x protein loading dye (SDS 3.4%).

## Data Availability

All data from this work are included in the main text and the SI Appendix of the paper.

## Acknowledgments

The work was funded by recurrent funds from CNRS and University of Bordeaux to S.J.S. and an ANR grant to S.J.S. (ANR SFAS AAP CE11). WD was supported in part by the National Science Centre of Poland (grant no. 2015/17/D/ST6/04054). We thank Robert Debuchy for providing the cosmids and plasmids for the cloning of *het-Q1*.

## Author contributions

C.C., S.J.S. and A.D. designed research; C.C., W.D., A.G.F., B.P. and A.D. performed research; C.C., W.D., S.J.S., and A.D. analyzed data; and A.D. wrote the paper

