## Supplementary Appendix for "Fungal gasdermin-like proteins are controlled by proteolytic cleavage"

**Corresponding authors**

Asen Daskalov

CNRS, UMR 5095, Institut de Biochimie et Génétique Cellulaires, University of Bordeaux, Bordeaux, France

Corinne Clavé

CNRS, UMR 5095, Institut de Biochimie et Génétique Cellulaires, University of Bordeaux, Bordeaux, France

**This PDF file includes:**

Supplementary Materials and Methods

Tables S1, S2, S3, S4

Figures S1 to S4

SI References

### Supplementary Materials and Methods

#### Sequence analysis and alignments

##### Homology search of *het-Q1*.

Homologs of three HET-Q1 proteins from *Podospora anserina* (XP_001908446.1, XP_001904890.1, XP_001910568.1) were searched amongst Ascomycota in the nr database (1) using PSI-BLAST (2) (75% overlap with the query, E-value up to 1-e20, 4 iterations). Out of 2072 non-redundant sequences found, 1854 with the length between 200 and 400 amino acids were selected for further analyses (2274 including identical sequences in different strains). Numbers of found *het-Q1* homologs and HeLo (PF14479) and Helo_like_N (PF17111) proteins reported by Pfam (4) were compared, in genomes with at least one occurrence of each, and their correlation was evaluated using the Pearson and Spearman correlation methods.

##### Identification of genomic neighbors of *het-Q1*.

An in-house Python script (aided by requests (Reitz, K.) and xmltodict (Blech) packages) was used to query NCBI (1) or EBI APIs (3) to fetch proteins coded by genes within the +/- 10 kbp neighborhood of the genes encoding 2274 *het-Q1* homologs. Due to various formatting issues of the database entries, neighborhoods were retrieved for 1884 het-Q1 homologs (83%) in 590 strains (401 species) identified with BioProject and BioSample accessions. The neighborhoods consisted of 12,380 protein-coding genes in total, including 1244 sequences annotated as Peptidase_S8 (PF00082) and 357 sequences annotated as CHAT (PF12770) using Pfam-A (4) at the sequence and domain E-values of 1e-2. Moreover, 178 neighborhoods included at least one other protease domain according to Pfam (E-value of 1e-2).

##### Annotation of Peptidase_S8- and CHAT-containing *het-Q1* neighbors.

The N- and C-termini of these proteins were extracted according to the Pfam-A hit envelopes. Next, 1440 N-termini and 591 C-termini at least 50 amino acids long were grouped using MMseqs2 (5) (clustering mode 1, pairwise sequence coverage of 80%, single-step clustering) into 547 clusters. For each cluster, sequences were aligned with Clustal Omega (6) (with the --auto option) and used as a query in HHblits (7, 8) search against Pfam-A (initial alignment trimmed at 50% gaps, E-value of 1e-3, 2 iterations). Eventually, Pfam-A annotations were assigned to N- and C-termini at least 100 amino acids long, according to hits in Pfam-A – direct and through HHblits search – at E-value of 1e-2 and to manual curation (aided with CDvist (9) and HHpred (10) web tools).

**Phylogenetic analysis of *het-Q1* homologs.**

The sequences of 1778 non-redundant *het-Q1* homologs were aligned using Clustal Omega (with the --auto option) and trimmed at 50% of gaps using trimAl (11). PhyML software (12) with default parameters was used to generate the maximum-likelihood phylogenetic tree. Branch support values were obtained in the approximate likelihood-ratio test using the Shimodaira-Hasegawa-like procedure (the aLRT SH-like method (13)). The tree was visualized using the ETE 3 package (14) in Python. Branches were collapsed at branch length (BRL) of 1.65.

**Protein extraction and Western blotting**

Total-protein extracts were prepared from strains grown on solid (SU) medium for 48h at 26°C. Protein extraction was realized in denaturing conditions (Urea 9M, SDS 1%, Tris HCl pH 6.8 25 mM, EDTA 1mM, β-mercaptoethanol 0.7 M) after freeze-drying of the samples. Electrophoresis (SDS-PAGE) was performed using standard procedures on 12% polyacrylamid gels. Proteins were transferred (2h at 40 mA/gel) to Amersham Hybond-ECL membrane (GE Healthcare) and immunoblotting was carried out with anti-V5 (ref. MCA1360 Biorad) antibodies in PBS-T buffer (PBS 1x, Tween-20 0.1% w/v).

**Fast protein liquid chromatography (FPLC)**

Mixtures of crude cell extracts were incubated for 30 min at room temperature (~22°C) and loaded directly on a Superose® 6 size-exclusion chromatography column, equilibrated with 10 mM PBS. Experiments were carried with an AKTA Purifier protein purification system at 4°C. FPLC fractionations (1 ml each) were kept on ice and subsequently analyzed by SDS-PAGE and immunoblotting.

**Table S1. Transformation efficiency of cosmid clones from the *PaATG1/het-Q1* region into *het-Q2* protoplasts**

| DNA | Total transformants |
|---|---|
| No DNA | 0 |
| Control plasmid pCSN43 | 77 |
| **CQ2** | |
| Transformation #1 | 50 |
| Transformation #2 | 38 |
| **CQ3** | |
| Transformation #1 | 77 |
| Transformation #2 | 68 |
| **CQ4** | |
| Transformation #1 | 180 |
| Transformation #2 | 44 |
| **CQ5** | |
| Transformation #1 | 27 |
| Transformation #2 | 34 |

**Table S2. Transformation efficiency of plasmid clones from the *PaATG1/het-Q1* region into *het-Q2* and *het-Q1* protoplasts**

| DNA | Transf. in *Q2* | Transf. in *Q1* |
|---|---|---|
| **No DNA** | 0 | 0 |
| **Control plasmid** | | |
| Transformation #1 | 125 | 73 |
| Transformation #2 | 60 | 62 |
| **Q1** | | |
| Transformation #1 | 161 | 29 |
| Transformation #2 | 263 | 34 |
| **Q2** | | |
| Transformation #1 | 47 | nd. |
| Transformation #2 | 42 | nd. |
| **Q3** | | |
| Transformation #1 | 34 | 23 |
| Transformation #2 | 45 | 20 |
| **Q4** | | |
| Transformation #1 | 5 | 25 |
| Transformation #2 | 5 | 31 |
| **Q5** | | |
| Transformation #1 | 15 | 11 |
| Transformation #2 | 9 | 18 |

**Table S3. Transformation efficiency of sub-clones of plasmid Q4 from the *PaATG1/het-Q1* region into *het-Q2* protoplasts**

| DNA | Genes present on construct | Transformants (plates counted) | Trns./plate |
|---|---|---|---|
| **No DNA** | - | 0 (10) | 0 |
| **Control plasmid pAN52.1GFP** | - | | |
| Transformation #1 | | 179 (6) | 29.8 |
| Transformation #2 | | 106 (6) | 17.6 |
| **Q4 plasmid** | Pa_7_10750, -60, -70, -75, -80 | | |
| Transformation #1 | | 5 (5) | 1 |
| Transformation #2 | | 8 (9) | 0.9 |
| **Q4 Δ*Bgl*II** | None | | |
| Transformation #1 | | 174 (5) | 34.8 |
| Transformation #2 | | 255 (7) | 36.4 |
| **Q4 Δ*Cla*I** | Pa_7_10750, -60, -80 | | |
| Transformation #1 | | 118 (8) | 14.7 |
| Transformation #2 | | 100 (6) | 16.6 |
| **Q4 Δ*Kpn*I** | Pa_7_10750, -75, -80 | | |
| Transformation #1 | | 22 (3) | 7.3 |
| Transformation #2 | | 29 (4) | 7.2 |
| **Q4 Δ*Nru*I** | Pa_7_10760, -70, -75, -80 | | |
| Transformation #1 | | 53 (9) | 5.9 |
| Transformation #2 | | 34 (7) | 4.8 |
| **Q4 Δ*Sac*I** | Pa_7_10780 | | |
| Transformation #1 | | 133 (8) | 16.6 |
| Transformation #2 | | 167 (7) | 23.8 |
| **Q4 Δ*Sal*I** | Pa_7_10775, -80 | | |
| Transformation #1 | | 17 (7) | 2.4 |
| Transformation #2 | | 16 (7) | 2.3 |
| **Q4 Δ*Xho*I** | Pa_7_10750, -60, -70 | | |
| Transformation #1 | | 170 (8) | 21.2 |
| Transformation #2 | | 98 (6) | 16.3 |

**Table S4. Transformation efficiency of plasmids bearing *het-Q1* and *het-Q2* into *het-Q1*, *het-Q2* and Δ*het-Q* protoplasts**

| | Total transformants | | |
|---|---|---|---|
| DNA | *het-Q1* recipient | *het-Q2* recipient | Δ*het-Q* recipient |
| **No DNA** | 0 | 0 | 0 |
| **Control plasmid SKhph-B3370** | | | |
| Transformation #1 | 366 | 128 | 232 |
| Transformation #2 | 166 | 261 | 301 |
| Mean | 266 | 194.5 | 266.5 |
| ***het-Q1*** | | | |
| Transformation #1 | 138 | 13 | 197 |
| Transformation #2 | 324 | 14 | 204 |
| Transformation #3 | 477 | 13 | 228 |
| Transformation #4 | 287 | 15 | 241 |
| Mean | 306.5 | 13.7 | 217.5 |
| ***het-Q2*** | | | |
| Transformation #1 | 10 | 172 | 340 |
| Transformation #2 | 21 | 220 | 225 |
| Transformation #3 | 11 | 304 | 277 |
| Transformation #4 | 24 | 243 | 310 |
| Mean | 16.5 | 234.7 | 288 |

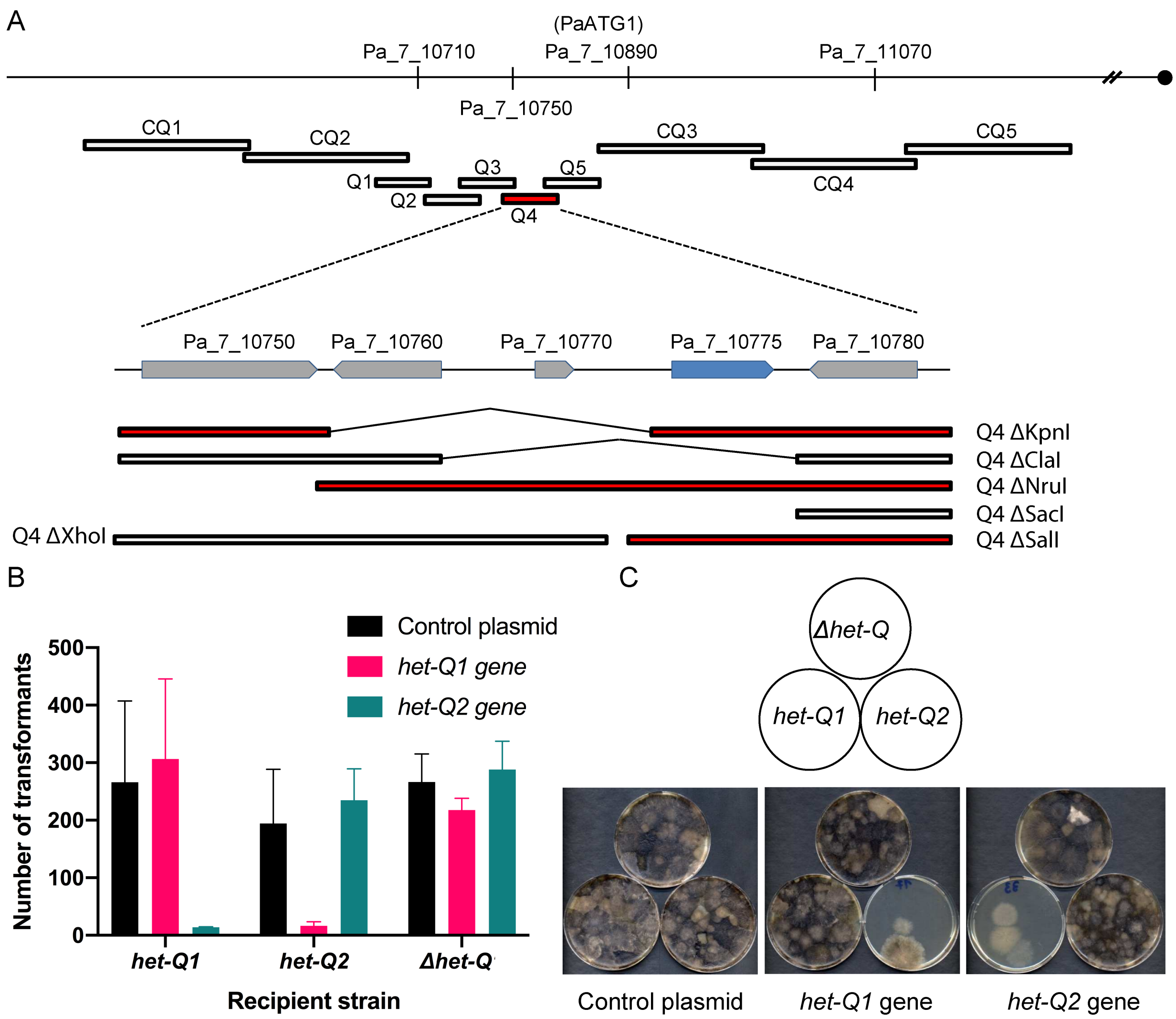


**Figure S1. Molecular identification and cloning of allorecognition cell death-inducing *het-Q1* and *het-Q2* genes from *P. anserina*. A.** Chromosome walk strategy to identify the *het-Q* locus, containing the idiomorphs *het-Q1* (gasdermin-encoding gene) and *het-Q2* (serine protease-encoding gene). The physical distance between the het-Q (Pa_7_10775) locus and PaATG1 (Pa_7_10890) is ~60 kbp. Cosmids (CQs) and plasmids (Qs) spanning the genomic region of the *het-Q1* locus, and found to reduce transformation viability of a *het-Q2* strain, are shown as red bars. Grey boxes indicate gene positions near the gasdermin-encoding *het-Q1* gene (blue box). **B.** Quantification of transformation viability of *P. anserina* strains from different *het-Q* genotypes. Strains are transformed with vectors carrying *het-Q1* or *het-Q2* and number of viable transformants are counted (Table S4). **C.** The *het-Q1* and *het-Q2* genes reduce strongly transformants viability when introduced in strains from the opposite genetic background. Cartoon (top) shows the genetic background for each recipient strain in the transformation plates shown below.

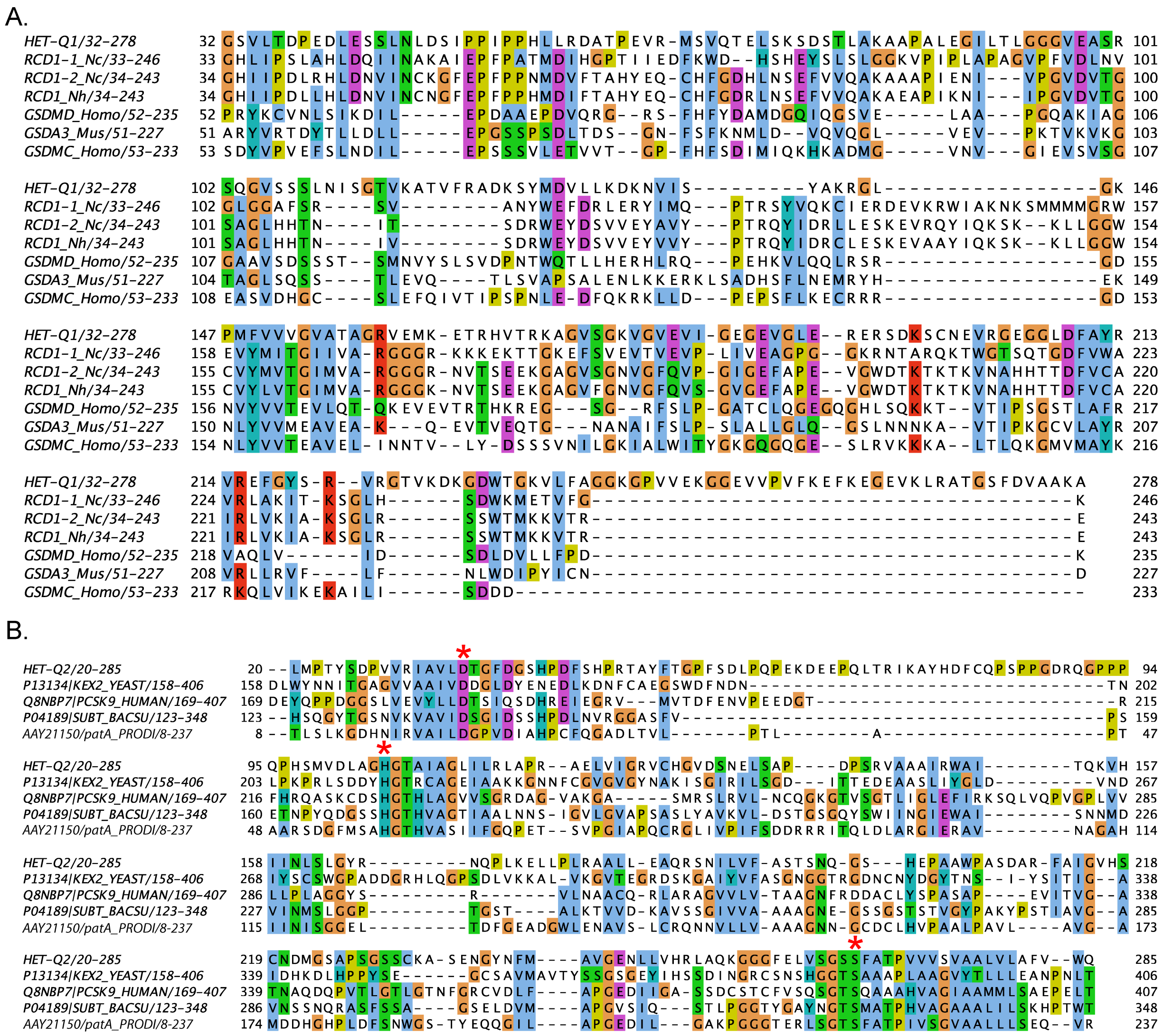


**Figure S2. Shown are protein sequence alignments of fungal and mammalian gasdermins (A) and of different subtilisin-like serine proteases (B). A.** Protein sequence of HET-Q1 from *P. anserina* is aligned with RCD-1 allelic variants from *Neurospora crassa* (RCD1-1_Nc (EAA33662.) and RCD1-2_Nc) and *Neurospora hispaniola* (RCD1_Nh) and the sequences of human gasdermin D (GSDMD_Homo, NP_079012), gasdermin C (GSDMC_Homo, NP_113603) and mouse gasdermin A3 (GSDMA3_Mus, NP_001007462). **B.** Protein sequence alignment of HET-Q2, yeast kexin protease (P13134), human subtilisin/kexin protease (Q8NBP7), subtilisin E (P04189) from *Bacillus subtilis* (bacteria) and a subtilisin-like protein (AAY21150) from *Prochloron didemni* (unicellular cyanobacteria). Amino acid residues defining the catalytic triad of these serine proteases are indicated with red asterisks. Conserved residues are shown in ClustalX colors, based on the nature of the amino acid residues: hydrophobic – blue, polar – green, negative charge – magenta, positive charge – red, aromatic – cyan, glycine – orange, proline – yellow, cysteine – pink, any/gap – white.

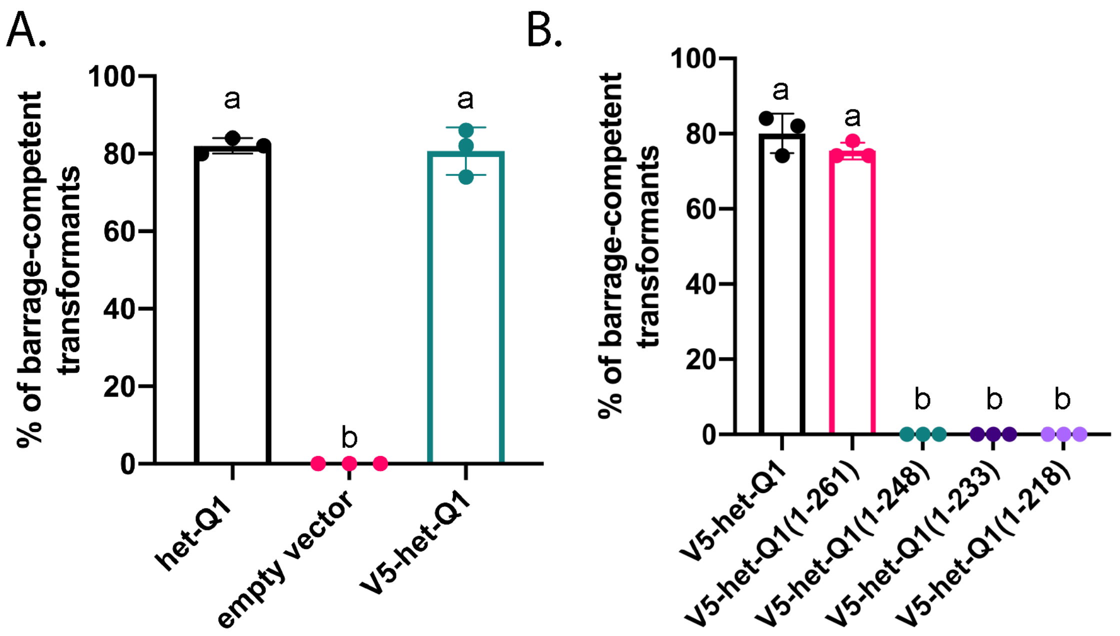


**Figure S3. Cell death-inducing ability of V5-tagged HET-Q1 constructs. A.** Transformants with a V5-tagged het-Q1 allele produce cell death at similar levels to non-tagged wild type het-Q1. Experiments were performed in triplicate with 50 transformants tested per experiment. P-value (a ≠ b) < 0.0001, one-way ANOVA with Tukey's multiple comparisons test. **B.** Quantification of barrage-inducing ability of truncated V5-tagged het-Q1 alleles. P-value (a ≠ b) < 0.0001, one-way ANOVA with Tukey's multiple comparisons test.

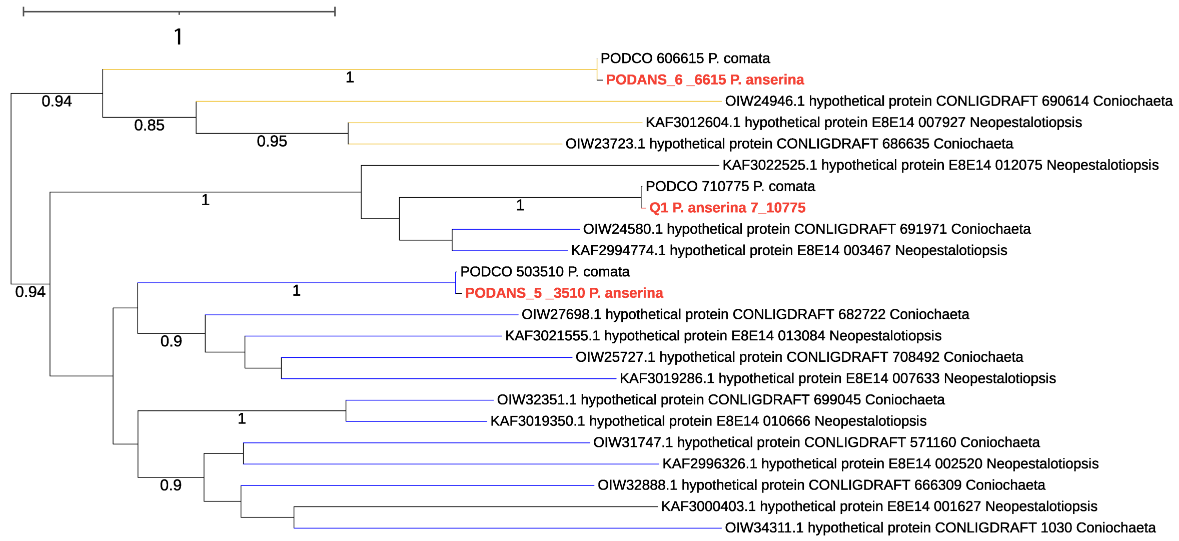


**Figure S4. Phylogenetic analyses of HET-Q1 gasdermins.** The three identified HET-Q1 homologs from *P. anserina* (Podans_7_10775, Podans_5_3510, Podans_6_6615) (coloured in red) are grouped by maximum-likelihood with HET-Q1 proteins from fungi. GeneBank ID is given for each protein. Branches are colored in accordance to the type of protease domain found on the HET-Q2-like proteins, encoded by the genes custered with the different *het-Q1* genes (blue – S8 family serine protease and yellow – caspase-like CHAT family, black – no identified *het-Q2-like* gene).
